# AreTomo: An integrated software package for automated marker-free, motion-corrected cryo-electron tomographic alignment and reconstruction

**DOI:** 10.1101/2022.02.15.480593

**Authors:** Shawn Zheng, Georg Wolff, Garrett Greenan, Zhen Chen, Frank G. A. Faas, Montserrat Bárcena, Abraham J. Koster, Yifan Cheng, David Agard

**Affiliations:** Dept. Biochemistry and Biophysics, University of California, San Francisco, USA; Howard Hughes Medical Institute; Section Electron Microscopy, Department of Cell and Chemical Biology, Leiden University Medical Center, Leiden 2333 ZC, Netherlands

**Keywords:** electron tomography, tomographic alignment, marker-free alignment, tomographic reconstruction, local beam-induced motion, GPU acceleration

## Abstract

AreTomo, an abbreviation for Alignment and Reconstruction for Electron Tomography, is a GPU accelerated software package that fully automates motion-corrected marker-free tomographic alignment and reconstruction in a single package. By correcting in-plane rotation, translation, and importantly, the local motion resulting from beam-induced motion from tilt to tilt, AreTomo can produce tomograms with sufficient accuracy to be directly used for subtomogram averaging. Another major application is the on-the-fly reconstruction of tomograms in parallel with tilt series collection to provide users with real-time feedback of sample quality allowing users to make any necessary adjustments of collection parameters. Here, the multiple alignment algorithms implemented in AreTomo are described and the local motions measured on a typical tilt series are analyzed. The residual local motion after correction for global motion was found in the range of ±80 Å, indicating that the accurate correction of local motion is critical for high-resolution cryo-electron tomography (cryoET).

## 1. Introduction

The combination of cryo-electron tomography (cryoET) with subtomogram averaging (STA) enables the *in-situ* study of proteins and macromolecular complexes at high resolutions (sub-nanometer) without purification. Important information is therefore preserved including the locations of proteins in their native environment and how they form, interact with, and are modulated by other macromolecular assemblies. The earliest STA application dates back to 1983 when negative-stained 30S ribosomal subunits of Escherichia coli were averaged (Knauer et al., 1983). Fourteen years later, the first cryoET based STA study was published (Walz et al., 1997). More recently, a new milestone was reached with STA-enabled high-resolution cryoET reaching subnanometer and even sub 5Å resolutions (Tegunov et al., 2021). Importantly, this approach can provide critical new insights into cell biology and mechanisms of pathogenesis. In one current example, the structures of the SARS-Cov-2 spike proteins were published with a map of the spike head at 7.9Å resolution and a map of the closed conformation at 4.9Å resolution (Turoňová et al., 2020). The characteristic hallmark of most in situ studies is the need for hundreds of tomograms to accumulate a sufficient number of subtomograms to enhance the high-resolution signal to noise ratio (SNR) and restore information lost due to the missing wedges. A major limitation, at least in the initial stages of STA is the information lost due to misalignment within each tomogram. This directly impacts the ability to identify and select the object of interest and perform the initial sub-volume alignments. Thus, while in good cases it is now possible to correct tomogram alignment errors by sub-volume polishing (see below), accurate starting tomographic alignment provides a critical foundation for high-resolution cryoET.

Although fiducial marker-based alignment has several drawbacks such as low throughput, almost inevitable manual intervention, and possible inconsistent movement of markers with respect to specimens, its precision has historically outperformed previous marker-free alignment efforts (Leigh et al., 2019). The 8.7Å reconstruction of the SARS-CoV-2 spike protein in pre- and post-fusion conformations is a typical example where 319 tomograms were generated based upon fiducial alignment (Yao et al., 2020). In separate research, the receptor binding domains (RBDs) of the spike surface protein were studied by subtomogram averaging using 340 tomograms collected and aligned using fiducial markers (Turoňová et al., 2020). While clearly this is a workable strategy, it is both labor intensive and the quality of the resultant alignment depends significantly on number and locations of the fiducial markers with respect to the area of interest.

However, the cryoET community has long recognized the value of developing marker-free tomographic alignment algorithms. While seeking ease and high throughput of the overall cryoET workflow are important driving forces, a more fundamental reason is that there are situations where adding fiducial markers, such as gold beads to the sample, is prohibitive for either technical or biological reasons, such as in cryo-lamella sample preparation (Rigort, 2012). There is also a concern that the use of colloidal gold may interfere with the sample (Han et al., 2014). The earliest marker free alignment for electron tomography was based upon cross-correlation of tilt images (Frank et al., 1992). Pairwise correlation between two images adjacent in tilt range was used to determine the translational shifts. Another early approach was based upon common lines (Liu et al., 1995). Since the common lines and the tilt axis have the same orientation, an iterative approach was developed to determine the orientations of the tilt axis as well as the translational shifts. More generally, projection matching, implemented in Protomo (Winkler et al., 2007), seeks to determine an optimal set of alignment parameters by iteratively matching equivalent specimen regions to a reference re-projection of an imperfectly reconstructed volume until no improvement can be made. It is computationally intensive since iterative calculation of alignment parameters requires repeated computation of back- and forward-projections. As a simplifying hybrid, feature-based alignment uses intrinsic sample features as virtual markers (Castaño-Díez et al, 2007; 2010). The movement of the features from tilt to tilt, called trails, are measured by cross correlation. Various metrics were developed to efficiently select the most useful trails from a pool of thousands of feature candidates. Another strategy for feature-based alignment selected a set of darkest voxels from the initial coarse-aligned tomogram to serve as 3D landmarks (Chen et al., 2019). These landmarks were then projected back to the tilt series and 2D patches centered at each projected landmark were extracted to generate a set of patch tilt series. The reconstructed local 3D volumes are used to refine the locations of the landmark. Using this strategy, subtomogram averaging of whole Escherichia coli overexpressing a double-layer-spanning membrane protein reportedly achieved 14Å resolution (Chen et al., 2019).

An important innovation to this approach was first achieved in emClarity which used the subtomograms themselves as fiducial markers (Himes et al., 2018). Analogous to particle polishing in single particle approaches but in 3D, the location of the subtomogram within each tilt in the raw tilt series were iteratively refined by the reference projections. The set of alignment parameters to be refined including shift, in-plane rotation, etc. were determined using overlapping patches, each of which contained a fixed number of particles. The reconstruction of the yeast 80S ribosome by emClarity reached 7.8Å. This has been more recently extended in several programs, i.e., M (Tegunov, 2021), Relion 4 (Kimanius, 2021), and BISECT/CSPT (Bouvette, 2021), to include local CTF corrections and potentially even higher order aberration corrections. While polishing strategies are extremely powerful, such approaches are contingent upon having high quality subtomogram averages. Ideally, much of the benefits of polishing should be obtainable directly from the raw tilt images.

Another advancing front in cryoET is the correction of beam-induced motion. In single-particle cryo-electron microscopy (cryoEM), the effect of beam-induced motion is limited to a single micrograph, allowing it to be corrected micrograph by micrograph. In cryoET, however, beam-induced motion has a sweeping effect throughout the entire tilt series, not only blurring each tilt image but also deforming samples gradually from tilt to tilt. Unless individual subtomogram-fiducials can be tracked throughout the raw tilt series, sample deformation can translate into errors in tilt series alignment. Fernandez et al. made an important innovation by starting with conventional fiducial marker-based alignment, and then using the residual alignment errors to estimate local sample deformation from tilt to tilt, modelled by polynomial surfaces (Fernandez et al., 2018) and later thin-plate splines (Fernandez et al., 2019). Subtomogram averaging was then carried out for purified T20S proteasomes in a thin sample (~15 nm thick) resulting in an improvement from 12.0Å without the correction of local sample motion to ~9.0Å. Unfortunately for a thicker basal body sample (~300 nm thick), only minimal improvement was observed (from 30.5Å to 29.0Å), indicating the problem is not yet solved.

Driven by the availability of instrumentation for FIB milling of cryo lamellae, there is now increasing focus obtaining on *in-situ* high-resolution cryoET by marker-free alignment. To meet this growing need, we developed the AreTomo package to provide a combination of rapid, fully automated marker free alignment, integrated reconstruction, and correction for local beam-induced tilt-to-tilt distortions. Thus, AreTomo provides an integrated marker-free solution to the need for high-throughput and full automation in a single GPU-accelerated software package.

Here we describe the mathematical background as well as the implementation of AreTomo. It endeavors to restore the information lost due to various sources of errors including translational misalignment, in-plane rotation, tilt-angle offset, and anisotropic local motion due to beam induced motion. AreTomo also extends the dose weighting scheme developed by Grant et al. (Grant et al., 2015) for cryoET by treating a tilt series as if it were a single movie containing all tilt images sorted in the order of acquisition. AreTomo has been successfully used for multiple projects including the tomographic alignment and reconstruction of the coronavirus molecular pore complex that likely passes RNAs across the membranes of the double-membrane vesicles (DMVs). This key component of the coronaviral replication organelles was analyzed in their native host cellular environment (Wolff et al., 2020). Because it is so low copy number, 33 cryo-lamellae and 53 highest quality tomograms were needed to obtain just ~600 suitable subvolumes. Automatic alignment and reconstruction with AreTomo greatly accelerated the entire process.

## 2. Method and results

The entire AreTomo workflow is fully automated and begins with loading a series of MRC-format motion-corrected tilt images and ends with saving either the aligned tilt series and/or the reconstructed tomogram also as MRC files. Users can choose either weighted back projection (WBP) (Radermacher, 1992) or simultaneous algebraic reconstruction technique (SART) (Anderson et al.,1984) to reconstruct their tomograms. AreTomo implements both global and local alignments. The global alignment determines the tilt angle offset, translations of the tilt images, and orientations of the tilt axis as it varies throughout the tilt series. While GPU acceleration significantly improves the throughput, the computation is made much more efficient by alternately refining the in-plane rotations and the translations until no further improvement can be made or the maximum number of iterations has been reached. Upon completion of the global alignment, local alignment can be started to correct for local motions due to the progressive sample deformation under repeated beam exposures. This is a computationally intense iterative process since local motions are measured at various locations throughout the tilt series. Our performance test showed that the global alignment reconstruction of a tilt series containing 61 K2 images could be done in roughly three minutes, while including local alignments required about twenty-one minutes. Local motion refinement is made optional to allow rapid real-time reconstruction especially for the on-line assessment of sample quality.

The datasets used here to demonstrate key AreTomo functionality are from a study of the coronavirus DMV-spanning molecular pore complex (Wolff et al., 2020) as well as from lamellae of arterivirus-infected cells. Data were collected at 300kV using a Titan Krios (Thermo Fisher Scientific) electron microscope equipped with a Gatan GIF Quantum energy filter and a Gatan K2 summit direct detection camera (Gatan, Pleasanton, CA). The energy filter slit width was 20 eV and the physical pixel size was 3.51Å. Individual movies each having 18 frames were collected in counting mode, every 2° or 3° covering a 100° − 120° range. The total dose deposited on the sample was 120 *e*^−^/Å^2^. More detailed information can be found in the supplementary material of Wolff et al. (2020). The collected movies were first corrected for beam-induced motion using MotionCor2 with 5×5 patches (Zheng et al., 2017). The corrected micrographs were then assembled into a single MRC file as the input to AreTomo. Based upon the reconstructed tomograms, the sample thickness was found to be in the range of 130-230 nm.

### 2.1 Determination of tilt-angle offset

In practice, because of beam-induced doming, the way the sample is milled and stage offsets, a zero-tilt setting of the tilt stage does not correspond to a sample normal to the incident beam. Because knowing the correct tilt angles is important for accurate calculation of alignment parameters, it is useful to determine a single tilt-angle offset and add that to the nominal readings from the microscope. In practice, tilt-angle offsets for non-milled samples are usually less than 4°, whereas they are often considerably larger for cryo-FIB prepared samples. Correcting the tilt-angle offset also effectively levels the specimen in the tomogram as shown in Fig. 1 for a case with an ~12° offset.

**Fig. 1.**
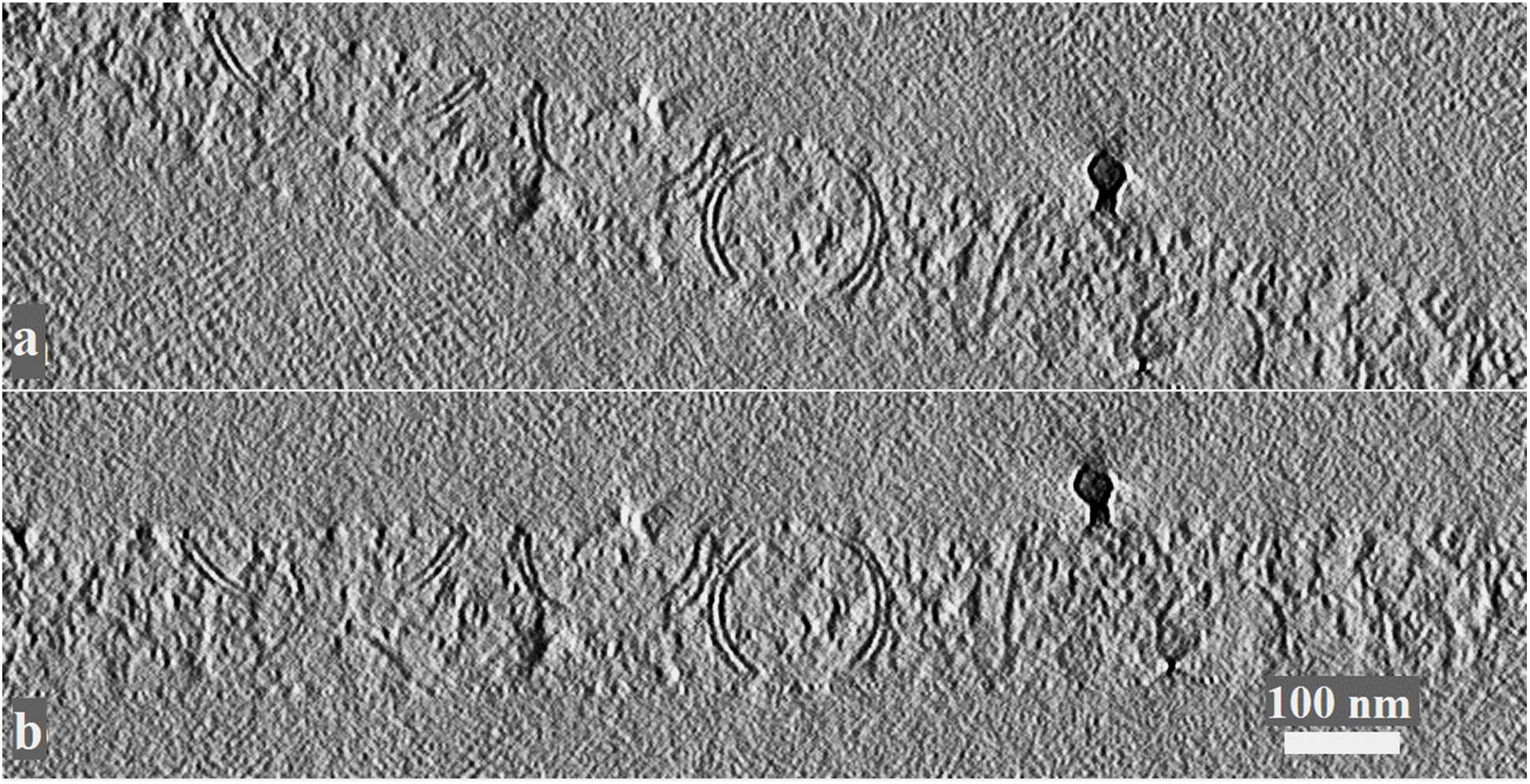
Central slices from *x-z* plane of a tomogram of DMVs in a coronavirus-induced cell (**a**) without and (**b**) with the correction of tilt-angle offset, respectively. The *x* axis is in the horizontal direction and perpendicular to the tilt axis. The *z* axis is in the vertical direction and parallel to the direction of electron beam. The measured tilt-angle offset is around −12°. The aligned tilt series was binned 6x by Fourier cropping for better visualization followed by the 3D reconstruction of weighted back-projection.

The effect of a large tilt-angle offset is shown in Fig 1(a) where the upper-left and the bottom-right structures were clipped. Correcting the tilt-angle offset allows a thinner, unclipped volume to be reconstructed more quickly. Importantly, the subsequent image alignment based on projection-matching benefits from the resultant thinner intermediate volume. This is because the voxels above and below the sample have non-zero values that bear no structural information yet when forward-projected, contribute noise that reduces the SNR of the computed projection images. As a result, both the efficiency and accuracy can be impaired if the tilt-angle offset is left uncorrected or if a more complex process to detect the sample extent (often called shrink-wrapping) is not implemented.

Beyond its impact on alignment, correcting the offset has several benefits. When a symmetric tilt range is used to collect a tilt series on a sample bearing a large tilt-angle offset, the actual tilting range will be skewed. Consequently, the sample can become excessively thick on one side of the tilt range, an inefficient way of using valuable electron dose. If the tilt-angle offset could be quickly measured in real time, subsequent tilt series could then be collected over an asymmetric nominal tilt range, cancelling out the estimated offset. A thinner, unclipped volume resulting from the correction of tilt-angle offset can be reconstructed more quickly, occupies less storage space and can be loaded into memory more rapidly for examination.

AreTomo seeks a tilt-angle offset, ∆*α*, that maximizes the sum of the cross-correlation coefficients (CC) of all pairs of adjacent tilt images. In each pair the image of higher tilt angle is stretched perpendicular to the tilt axis according to Eq. (1). The object function is given in Eq. (2) where *α*_*i*_ and *α*_*j*_ are tilt angles of the tilt images *t*_*i*_ and *t*_*j*_ in the same pair, respectively. *t*_*j*_ denotes the image at the higher tilt angle.

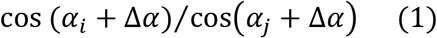

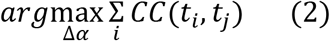

### 2.2 Alignment of in-plane rotations

In single-axis tomography, projection images intersect each other in Fourier space on common lines. Since the common lines have the same orientation as the tilt axis, they were first used by Liu et al. to determine the in-plane rotation for the freely supported specimen by maximizing the cross correlation of the common lines (Liu et al., 1995). Owen et al. introduced projection matching to refine the rotational alignment determined by the common line method (Owen et al., 1996). AreTomo adopted this latter approach but with some notable changes. Since the alignments for translation and in-plane rotation are inter-dependent, they are staggered in iterations, i.e., the rotational alignment is based upon the aligned tilt series from the previous iteration and the improved estimates of tilt axis orientations are applied for the next round of the translational alignment. The missing areas due to the correction of both translational and rotational misalignment are excluded in the common line calculation. Although the common line is theoretically the 1D projection of the 2D image in the direction perpendicular to the tilt axis, the 1D projection of the entire image would result in an inaccurate estimate of the common line since tilting increases the field of view of the sample, bringing in extra information that is absent in the lower tilt images. To suppress this source of error, AreTomo first determines the common fields of view throughout the aligned tilt series over which the common lines are calculated. The common fields are usually not rectangular due to the correction of the translation and in-plane rotation. The non-empty subarea of the zero-tilt image is used as the geometric reference for the common fields at other tilt angles. Lastly, realizing that while the tilt axis orientation varies from tilt to tilt, it does so slowly, a third-order polynomial function of tilt angles is used to model the variation while suppressing noise and solved using the conjugate gradient method by maximizing the following target function. The solution is a set of polynomial coefficients, *a*_0_, … , *a*_3_, that maximizes the sum of the correlation coefficient, *CC* in Eq. (3), of each pair of the common lines *l*_*i*_ and *l*_*j*_ where *i* and *j* are the indices of the tilt images.

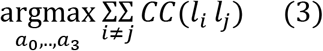

The better estimates of the tilt axes are then used to refine the translational alignment in the next iteration, which can be used again to improve the determination of the common fields and the tilt axes.

### 2.3 Translational alignment

The translational alignment implemented in AreTomo is a variant of the projection matching method. Central to the changes made in AreTomo is the weighting scheme used in the calculation of the projection images. According to the central section theorem, the Fourier transform of a 2D projection of an object is a slice of the 3D Fourier transform of that object. In Fourier space, an adjacent pair of slices bears the most resemblance whereas any two slices that are 90° apart are independent from each other except along the common lines. More precisely, the actual coupling between tilts is a function of resolution and Z thickness as well as the tilt angle difference. However, for generality, when the intermediate tomogram is reconstructed for the subsequent forward projection to a reference angle, AreTomo weights the tilt images based upon the cosine of the angular differences relative to the reference angle. When the angular difference exceeds 90°, its supplementary angle is used instead. The tilt image collected at the reference angle is excluded to avoid self-correlation. Another change is tilt images are aligned sequentially from low- to high-tilt angle with the image having a corrected tilt angle closest to zero set to be the reference. Only the aligned tilt images plus the reference are used to reconstruct the intermediate tomogram, which is then forward-projected to the angle of the next tilt image to be aligned. As such, more images are used for the alignment of images at higher angles. This approach is less prone to the accumulation of errors compared to using only a pairwise comparison. It should be noted that, although the entire alignment process is iterative, the actual calculation of the translational offset is non-iterative, using a filtered FFT-based cross correlation for efficiency.

### 2.4 Local alignment

Local alignment in AreTomo is founded on a series of local measurements that track local features distributed throughout the field of view over the entire tilt range. Instead of picking a small set of reference points such as gold beads (Fernandez et al., 2018, 2019) or manually picking features, the zero-tilt image is subdivided into patches that are tracked throughout the tilt series. Noting that the patches are the projections of subvolumes, tracking patches is therefore equivalent to tracking in the image plane the movements of subvolumes from tilt to tilt. Since the tilt series has been globally aligned, the projected centers of each subvolume at all other tilt angles can be pre-estimated based upon geometric rotation when the sample thickness is ignored. In each local measurement, instead of actually subdividing the whole tilt series, the estimated subvolume center at each tilt is shifted to the center of the field of view by applying an induced shift to the corresponding tilt image followed by applying a soft mask. Projection matching alignment is then applied to the shifted and masked tilt series to yield a series of residual shifts, one per each tilt for each subvolume. Summing each induced shift and the corresponding residual shift produces a path that can be expressed as a set of coordinates (*u*_*ij*_, *v*_*ij*_, *α*_*j*_) where *i* and *j* specify the *i*th subvolume and the *j*th tilt angle respectively. *u* and *v* are the coordinates in horizontal and vertical axes, respectively in the image plane. *α* denotes the tilt angle. The hypothesis is that the paths are the joint effect of geometric rotation resulting from sample tilting and local deformation due to beam induced motion. To separate these two movements, the paths are fit to a 3D model of rigid-body rotation. The residuals of (*u*, *v*) deviating from the model are treated as the local deformation whereas the projected centers of the subvolumes at different tilt angles are given by the model. A major challenge in marker-free local alignment is that, although beam induced motion is limited within the neighborhood of the subvolumes, sample tilting can shift them far away – even after the tilt series is globally aligned. The combined movement of a subvolume from tilt to tilt mainly depends upon how far it is from the tilt axis, which is unknown since its depth inside the sample is unknown. AreTomo implements a two-step tracking scheme. Initially, since the path of each subvolume is initially coarsely estimated without taking into account sample thickness, a soft mask covering the entire field of view is applied to the tilt series before the translational alignment is performed. The measured translations are then used to update the coordinates of the estimated path. The updated path, although more truthfully representing the subvolume movement, still bears errors resulting from remote signals not eliminated by the large mask. A much tighter soft mask whose diameter is one eighth of the whole field of view is therefore applied in the second step to better resolve the local motion. This process is repeated for the remaining subvolumes.

For a tilt series of a well-dispersed sample, users can simply specify in the command line the number of patches along each image axis. For a sparse sample, users can provide a list of coordinates representing the locations where the local alignments will be performed. This is a guided approach to avoid erroneous alignment carried out on the empty areas. Regardless of which approach is taken, a sanity check is performed at the end of each cycle to identify any local measurements having abnormally large magnitudes compared to their neighbors within the same tilt image. The outliers are excluded in the subsequent correction for the local motion. Since the local motion is measured at a limited number of discrete locations, a distance-weighted interpolation scheme is then used to smoothly correct local motion at each pixel. Let *U*(*x*, *y*) and *V*(*x*, *y*) be the horizontal and vertical components, respectively, of the interpolated local motion at pixel (*x*, *y*) calculated according the following equations.

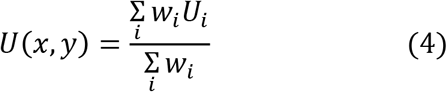

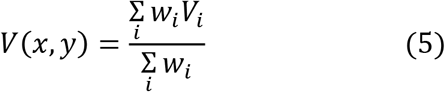

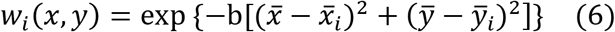

In Eqs. (4) and (5), *U*_*i*_ and *V*_*i*_ are the measured local motions of the *i*th patch. *w*_*i*_ represents the contribution of the local motion from the *i*th patch to pixel (*x*, *y*). In Eq. (6) b is a damping factor that controls the outreach of the measured local motions. (*x*_*i*_, *y*_*i*_) are the coordinates of the *i*th patch. The overbar indicates the coordinate is normalized. Note that the correction for the local motion is integrated with the global correction. As a result, the raw images are interpolated only once to generate the aligned tilt series.

At this pixel spacing, the local motions measured among the testing data sets were typically less than 20 pixels. This is probably why we observed that the smaller DMVs in local-aligned tomograms often show more noticeable improvement than the larger ones. Hence, we used a tomogram reconstructed from a tilt series collected on DMVs in arterivirus-infected cells to demonstrate the improvement since they are typically much smaller than the DMVs in coronavirus-infected cells. Fig. 2 presents a slice from the *x-z* plane of the tomogram reconstructed with both global (Fig 2a) and local alignment (Fig 2b). Since the specimens were ubiquitous in the field of view, the local alignment was performed by tracking the features in 36 patches that evenly divided the raw image at 0° of nominal reading in its *x* and *y* directions. For visualization here, the aligned tilt series was binned 6x by Fourier cropping followed by the 3D reconstruction of weighted back-projection. Three pairs of DMVs were chosen as examples to highlight the improvement resulting from the local alignment correction. The corresponding pairs are highlighted in the boxes of the same color. As can be seen, the local-alignment based DMVs are more spherical compared to their more peach-like counterparts in Fig. 2(a), a sign of the improved alignment since DMVs are generally spherical.

**Fig. 2.**
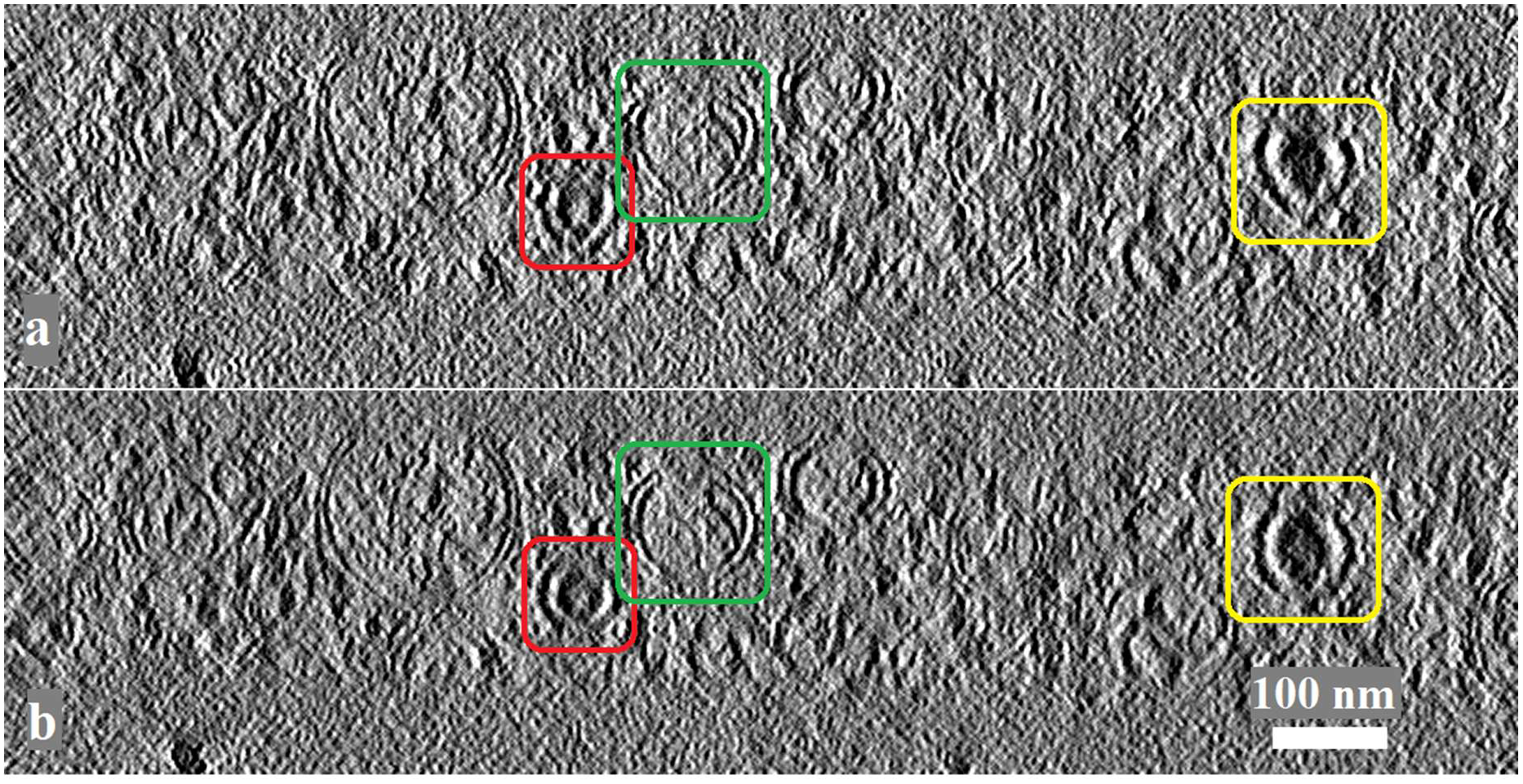
(**a**) A tomographic slice from an *x-z* plane reconstructed with only the global alignment. (**b**) The corresponding *x-z* slice reconstructed with both global and local alignments. The local alignment was performed by tracking the underlying features in 36 patches. The corresponding DMV pairs in arterivirus-infected cells are highlighted in equivalently colored boxes.

A tomographic slice from an *x*-*y* plane is given in Fig. 3 as an example of coronavirus-induced DMVs aligned and reconstructed by AreTomo. The entire processing was fully automated. To better visualize the molecular pore complex that spans the double membrane of one of the DMVs, Fig. 3 shows only the 6x binned tomogram reconstructed with the global and local alignments. The DMV-spanning molecular pore is highlighted in the white box. The zoom-in view is given in the inset with the molecular pore highlighted inside the red circle.

**Fig. 3.**
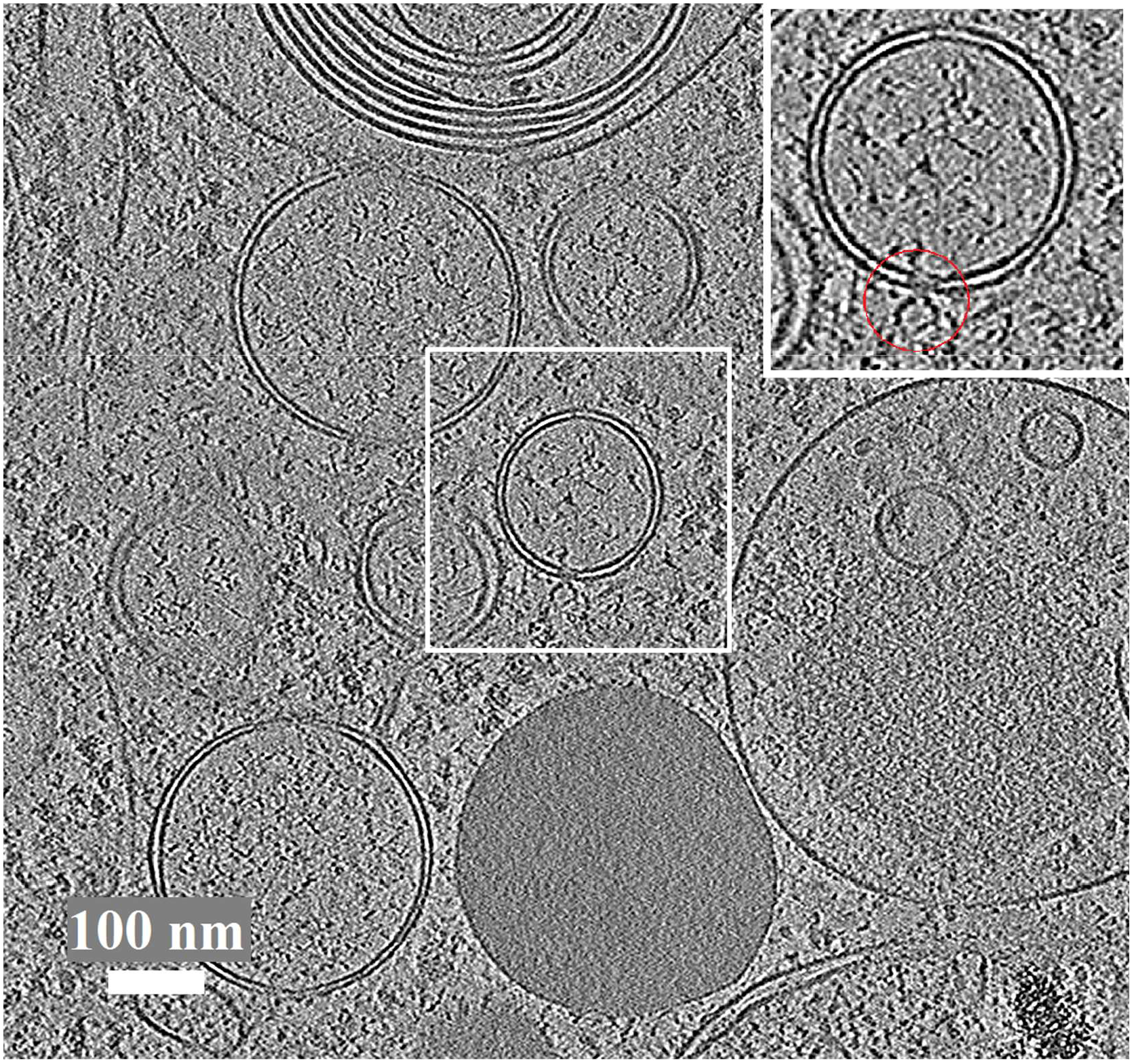
A tomographic slice from the *x-y* plane reconstructed from a tilt series both global- and local-aligned. The local alignment was based upon tracking 36 features. The aligned tilt series was binned 6x by Fourier cropping and then reconstructed by weighted back-projection.

Although central to our effort was the development of a robust tomographic alignment scheme, we are also interested in how the local motion varies over the field of view and from tilt to tilt. Such information can help develop a better alignment strategy and perhaps guide the data collection. Fig. 4 plots the distribution of local motions over the field of view of four tilt images acquired at −31°, 31°, −51°, and 51°, respectively. The vectors were magnified 20x to enhance their visibility. The green dots represent the modelled locations of the tracked features based only on the global motion correction, while the vector tips represent the measured locations including local motions. The tilt axis is vertical and at the center of the field of view. Our results show that the magnitude of the local motion can be as much as 80Å, while the differential magnitude across the image can be twice that. Such severe local motion, if left uncorrected, can be devastating to high-resolution cryoET. Correcting as much local motion as possible at the offset, rather than only after sub-volume refinement, should improve the quality of the initial subvolume alignment and directly facilitate the bootstrap process of iterative sub-volume refinement. At the very least it should make the entire non-linear process more robust and efficient.

**Fig. 4.**
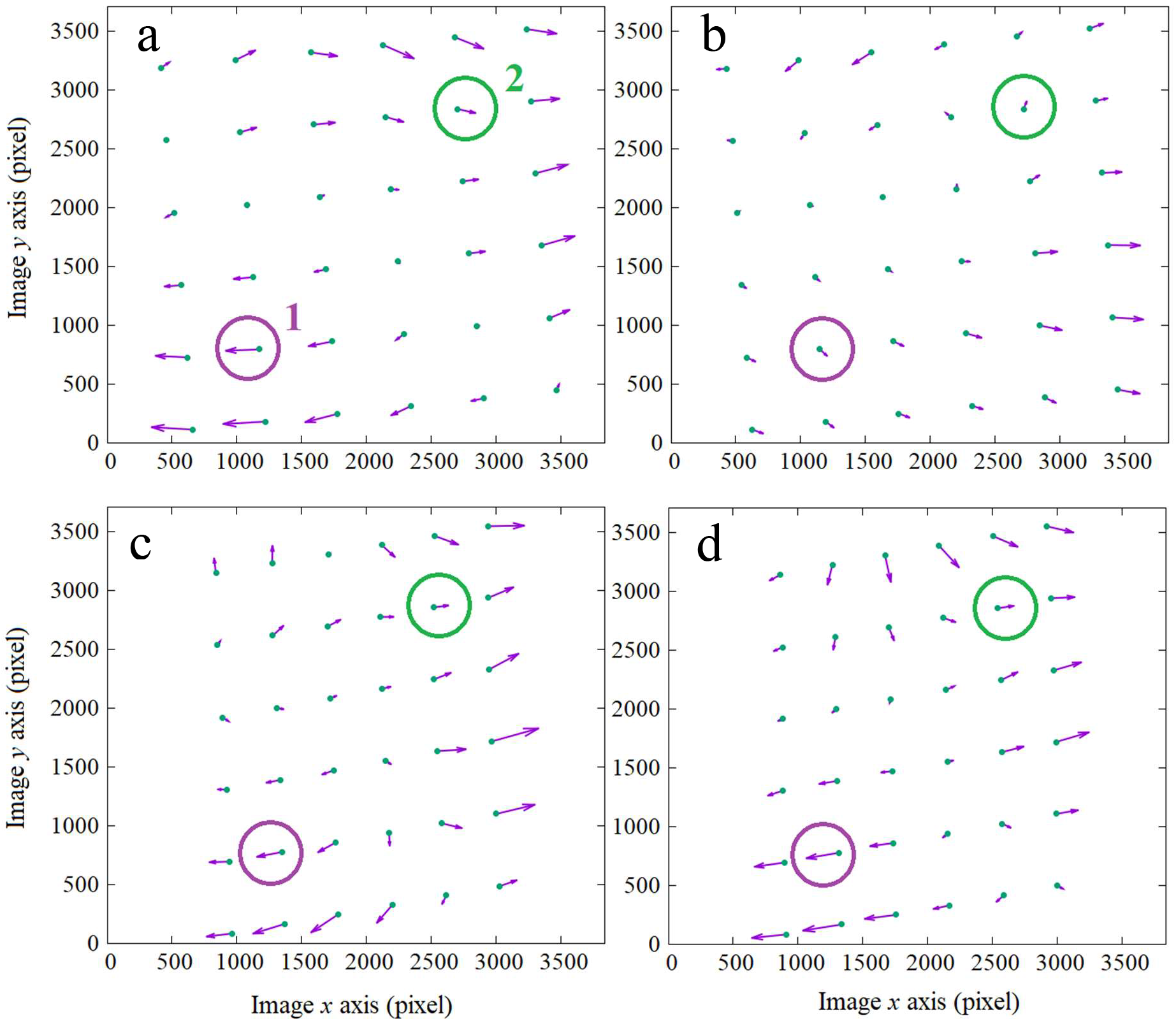
Vector distributions of measured local motions over the fields of views of four tilt images at (**a**) −31°, (**b**) 31°, (**c**) −51°, and (**d**) 51°, respectively. The vectors are magnified 20x to enhance the visibility. The green dots denote the modelled locations of the features if there were no local motion. The vector tips represent the actual locations. Two features were chosen to show their local motions over the entire tilt range in **Fig. 5**. The purple and green circles are used to label the local motions of these two features in all the subplots.

A region populated with smaller vectors can be observed in the upper-left quadrant in all subplots of Fig. 4 and likely corresponds to the doming center according to the doming model (Brilot et al., 2012; Zheng et al., 2017). To illustrate how their local motions change over the entire tilt range, two patches with their motion vectors labeled inside the circles in Fig. 4 were intentionally chosen from the diagonal sides of the doming center. The horizontal components of these vectors, i.e., the motion perpendicular to the tilt axis, were plotted against the tilt angle and are presented in Fig. 5. According the doming geometry, a component perpendicular to the tilt axis is the projection of the out-of-plane sample motion in the image plane.

**Fig. 5.**
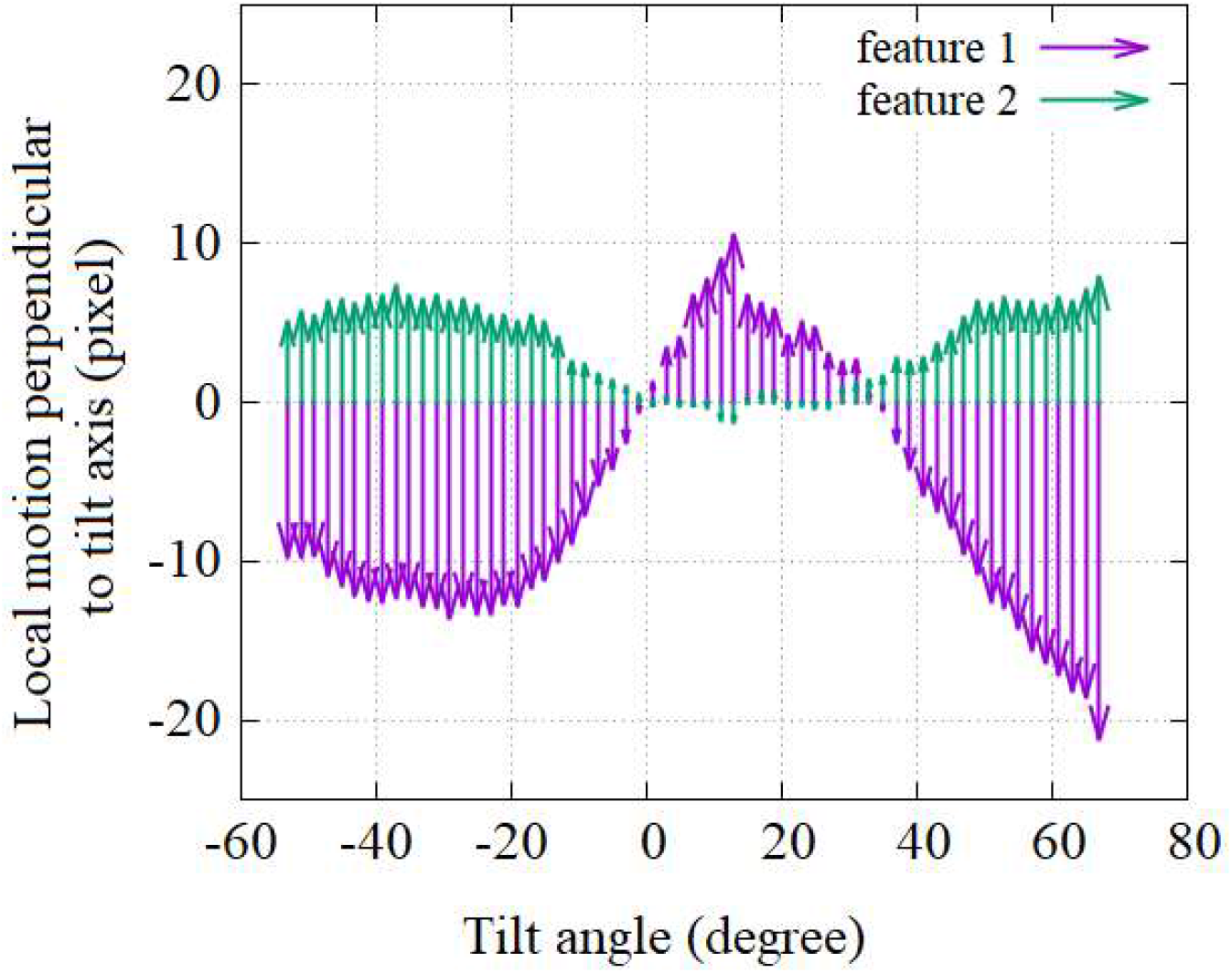
Variations of the local motions with respect to the tilt angles. The vectors are the components perpendicular to the tilt axis and therefore the projections of the out-of-plane sample motion. The corresponding features were selected from the diagonal sides of the doming center as shown in **Fig. 4**.

These two distributions are approximately anti-symmetric, in particular, at higher tilt angles. Since these two features were on the diagonal sides of the doming center, such an approximate symmetry is also consistent with the tilted doming model (Zheng et al., 2017). We can also see that the maximum doming motion is as much as 25 pixels (purple region), or equivalently 88 Å, given the pixel size of 3.51 Å. This in turn corresponds to an out-of-plane Z motion in excess of 100Å. Since a cryo sample domes during the exposure regardless of the tilt angle, it is likely that the out-of-plane motion can be equally severe when a sample is not tilted. This suggests that the defocus also varies for at least an equal amount from the first to the last movie frame in a single-particle cryoEM image. Such a relatively large defocus variation has implications for the current motion-correction strategies that simply add the motion-corrected frames together to form the final micrograph without taking into account these local defocus changes within the movies. Though the contribution of such defocus variation to the final reconstruction is unclear, further exploring this effect is worthwhile. It is also worth noting that since the local alignment can yield the z distribution of the underlying features, it is possible to take into account the z offset for 3D CTF correction when subtomograms are generated by means of local reconstructions instead of being extracted from the whole volume.

### 2.5 Performance

The performance test was carried out on a tilt series containing 61 tilt images of 3838 × 3710 pixels. It took 336 seconds to generate a 2x binned tomogram of 1918 × 1854 × 600 voxels with only the global alignment on a Linux system using a single NVIDIA Titan X GPU card that has 12 GB of RAM. When the local alignment was turned on with the underlying features in 36 patches (6 × 6) being tracked, the time went up to 4857 seconds to generate the same-size tomogram on the same system. On another Linux system using a single NVIDIA GV100 GPU card that has 32 GB of RAM, the same test took 156 seconds and 1618 seconds, respectively.

## 3 Conclusion

In principle, fractionating sample dose into a tomographic data collection should add additional information to aid in determining particle orientation and classification. Yet in practice cryoET has lagged well behind single particle methods in terms of resolution and robustness, even for equivalently thin samples. While many factors are responsible, perhaps the most important are the low signal to noise of each tilt, lack of coherence across tilts due to differential beam-induced motion, and the challenge of collecting and processing hundreds let alone thousands of tomograms. AreTomo aims to help by enabling a fully automated and efficient workflow that can go from a raw tilt series to a final tomogram without any manual intervention and its ability to correct of local beam induced motions throughout the entire tomogram. The local motions are of sufficient magnitude (>80Å) that, left uncorrected, they would dramatically degrade the quality of the tomogram. This continues the philosophy begun in our beam-induced motion correction software (MotionCor2) of doing as much polishing (in this case subvolume polishing) as possible in pre-processing, such that all subsequent steps are optimized.

It has been shown that AreTomo can generate tomograms with sufficient alignment accuracy that they can directly be used for subtomogram averaging. It is worth noting that AreTomo can generate a global-alignment based tomogram much faster than the collection of a tilt series, making it possible to reconstruct tomograms on the fly. While there is a long to-do list for AreTomo, we believe the current implementation is a valuable addition to the cryoET toolkit. This belief motivated us to publish the methodology and algorithms developed for AreTomo to promote the interest in the development of better marker-free tomographic software packages.

AreTomo can be downloaded from https://msg.ucsf.edu/software and is free for academic use.

## Acknowledgment

The authors are thankful to Dr. J. Lefman and G. Thomas-Collignon for support and discussion of code optimization. NVIDIA Corporate kindly and generously provided two NVIDIA Quadro GP100 cards and two NVIDIA Quadro GV100 cards. Y.C. is an investigator of Howard Hughes Medical Institute. This work was supported by NIH grants R35GM118099 (DAA) and 1R35GM140847 (YC) as well as NIH S10 equipment grants 1S10OD026881, 1S10OD020054, and 1S10OD021741. We thank David Bulkley, Glenn Gilbert and Matt Harington for their invaluable support of the UCSF cryoEM facility.

The authors are grateful to Prof. Eric Snijder for his critical support of our virology research at Leiden University Medical Center. The data on virus-infected samples were collected at the Netherlands Centre for Electron Nanoscopy (NeCEN) made possible through financial support from the Dutch Roadmap Grant NEMI (NWO grant 184.034.014).

---

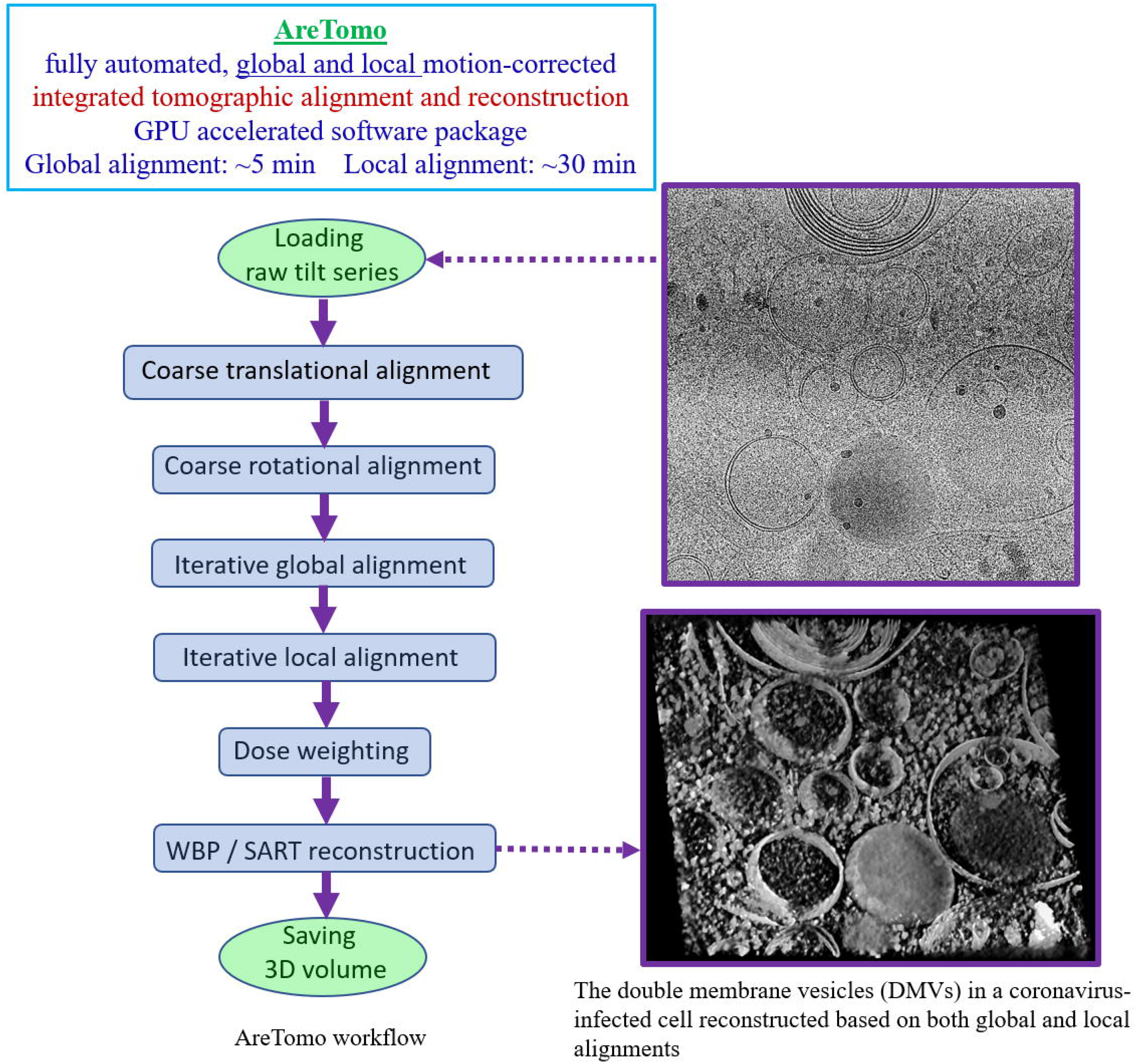

## References

Andersen, A., Kak, A., 1984. Simultaneous algebraic reconstruction technique (SART): a superior implementation of ART. Ultrasonic Imaging 6 (1), 81–94.

Bharat, T.A.M., Kureisaite-Ciziene, D., Hardy, G.G., Yu, E.W., Devant, J.M., Hagen W.J.H., Brun, Y.V., Briggs J.A.G., Lowe, J., 2017. Structure of the hexagonal surface layer on Caulobacter crescentus cells. Nat. Microbiol 2,17059

Bouvette, J., Liu, H., Du, X., Zhou, Y., Sikkema, A.P., Juliana da Fonseca Rezende e Mello, Klemm, B.P., Huang, R., Schaaper, R.M., Borgnia, M.J., Bartesaghi, A., 2021. Beam image-shift accelerated data acquisition for near-atomic resolution single-particle cryo-electron tomography. Nat. Communication (2021) 12:1957

Brilot, A.F., Chen, J.Z., Cheng, A., Pan, J., Harrison, S.C., Potter, C.S., Carragher, B., Henderson, R., Grigorieff, N., 2012. Beam-induced motion of vitrified specimen on holey carbon film. J. of Struct. Biol. 177, 630–637.

Castaño-Díez, D., Al-Amoudi, A., Glynn, A.M., Seybert, A., Frangakis, A.S., 2007. Fiducial-less alignment of cryo-sections. J. Struct. Biol. 159, 413–423.

Castaño-Díez, D., Scheffer, M., Al-Amoudi, A, Achilleas S. Frangakis A.S., 2010. Alignator: A GPU powered software package for robust fiducial-less alignment of cryo tilt-series. J. Struct. Biol. 170, 117–126.

Chen, M., Bell, J.M., Shi, X., Sun, S.Y., Wang, Z., Ludtke, S.J., 2019. A complete data processing workflow for cryo-ET and subtomogram averaging. Nat. Methods 16, 1161–1168.

Fernandeza, J-J., Li, S., Tanmay A.M. Bharat, T.A.M., Agard, D.A., 2018. Cryo-tomography tilt-series alignment with consideration of the beam-induced sample motion. J. Struct. Biol. 202, 200–209.

Fernandeza, J-J., Li, S., Agard, D.A., 2018. Consideration of sample motion in cryo-electron tomography based upon alignment residual interpolation. J. Struct. Biol. 205, 1–6.

Forster, F., Hegerl, R., 2007. Structure determination in situ by averaging of tomograms. Method Cell Biol 79, 741–767.

Frank, J., McEwen, B.F., 1992. Alignment by cross-correlation. Electron Tomography. Springer 205–213.

Grant, T., Grigorieff, N., 2015. Measuring the optimal exposure for single particle cryo-EM using a 2.6 A reconstruction of rotavirus VP6. eLife 4, 1–19

Han, R., Zhang, F., Wan, X., Fernández, J-J., Sun, F., Liu, Z., 2014. A marker-free automatic alignment method based on scale-invariant features. J. Struct. Biol. 186, 167–180. http://dx.doi.org/10.1016/j.jsb.2014.02.011

Himes, B.A., Zhang, P., 2018. emClarity: software for high-resolution cryo-electron tomography and subtomogram averaging. Nat. Methods 15, 955–961.

Kimanius, D., Dong, L., Sharov, G., Nakane, T., Scheres, H.W.S., 2021. New tools for automated cryo-EM single-particle analysis in RELION-4.0. bioRxiv, 2021.09.30.462538, doi: https://doi.org/10.1101/2021.09.30.462538

Knauer, V., Hegerl, R., Hoppe, W., 1983. Three-dimensional reconstruction and averaging of 30 S ribosomal subunits of Escherichia coli from electron micrographs. J. Mol. Biol. 163, 409–430.

Leigh, K.E., Navarroc, P.P., Scaramuzzac, S., Chen, W., Zhang, Y., Castaño-Díez, D., Kudryashev, M., 2019. Subtomogram averaging from cryo-electron tomograms. Method in Cell Biol. 152, 217–259. https://doi.org/10.1016/bs.mcb.2019.04.003

Liu, Y., Penczek, P. A., McEwen, B. F., Frank, J., 1995. A marker-free alignment method for electron tomography. Ultramicroscopy 58, 393–402.

Owen, C.H., Landis, W.J., 1996. Alignment of electron tomographic series by correlation without the use of gold particles. Ultramicroscopy 63, 27–28

Radermacher, M., 1992. Weighted back-projection methods. Electron Tomography, Springer 91–115. Rigort, A., Bauerlein, F.J.B., Villa, E., Eibauer, M., Laugks, T., Baumeister, W., Plitzko, J.M., 2012. Focused ion beam micromachining of eukaryotic cells for cryoelectron tomography, PNAS 109, 4449–4454

Schur, F. K.M., 2019. Toward high-resolution in situ structural biology with cryo-electron tomography and subtomogram averaging. Cur. Opinion in Struct. Biol. 58, 1–9

Tegunov, D., Xue, L., Dienemann, C., Cramer, P., Mahamid, J., 2021. Multi-particle cryo-EM refinement with M visualizes ribosome-antibiotic complex at 3.5Å in cells. Nat. Method 18, 186–193

Turoňová, B., Sikora, M., Schürmann, C., Hagen, W.J.H, Welsch, S., Blanc, F.E.C., Bülow, S., Gecht, M., Bagola, K., Hörner, C., Zandbergen, G., Landry, J., Azevedo N.T.D., Mosalaganti, S., Schwarz, A., Covino, R., Mühlebach, M.D., Hummer, G., Locker, J.K., Beck, M., 2020. In situ structural analysis of SARS-CoV-2 spike reveals flexibility mediated by three hinges. Science 370, 203–208

Walz, J., Tamura, T., Tamura, N., Grimm, R., Wolfgang Baumeister, W., Koster, A.J., 1997. Tricorn Protease Exists as an Icosahedral Supermolecule In Vivo. Molecular Cells 1, 59–65

Winkler, H., Zhu, P., Liu, J., Ye, F., Roux, K.H., Taylor, K.A., 2009. Tomographic subvolume alignment and subvolume classification applied to myosin V and SIV envelope spikes. J. Struct. Biol. 165, 64–77

Yao, H., Song, Y., Chen, Y., Wu, N., Xu, J., Sun, C., Zhang, J., Weng, T., Zhang, Z., Wu, Z., Cheng, L., Shi, D., Lu, X., Lei, J., Crispin, M., Shi, Y., Li, L., Li, S., 2020. Molecular Architecture of the SARS-CoV-2 Virus. Cell 183, 730–738.

Zhang, P., 2019. Advances in cryo-electron tomography and subtomogram averaging and classification. Current Opinion in Struct. Biol. 58, 249–258

Zheng, S.Q., Palovcak, E., Armache, J.P., Verba, K.A., Cheng, Y., Agard, D.A., 2017. Motioncor2: Anisotropic correction of beam-induced motion for improved cryo-electron microscopy. Nature Methods 14, 331–332.

